# Longitudinal changes in functional connectivity networks in the first year following stroke

**DOI:** 10.1101/2025.03.10.642404

**Authors:** Y. Tao, T.T. Schnur, J.H. Ding, R. Martin, B. Rapp

**Affiliations:** Department of Cognitive Science, Johns Hopkins University, Baltimore, Maryland, USA; Physical Medicine and Rehabilitation, University of Texas Health Houston, Houston, Texas, USA; State Key Laboratory of Cognitive Science and Mental Health, Institute of Psychology, Chinese Academy of Sciences, Beijing, China; Psychological Sciences, Rice University, Houston, Texas, USA; Department of Psychological and Brain Sciences, Johns Hopkins University, Baltimore, Maryland, USA; Department of Neuroscience, Johns Hopkins University, Baltimore, Maryland, USA; Department of Neurology. Johns Hopkins University, Baltimore, Maryland, USA

## Abstract

The functional organization of the brain consists of multiple subsystems, or modules, with dense functional communication within modules (i.e., visual, attention) and relatively sparse but vital communications between them. The two hemispheres also have strong functional communications, which presumably supports hemispheric lateralization and specialization. Subsequent to stroke, the functional organization undergoes neuroplastic changes over time. However, empirical longitudinal studies of human subjects are lacking. Here we analyzed three large-scale, whole-brain resting-state functional MRI connectivity measures: *modularity*, *hemispheric symmetry* (based on *system segregation*), and *homotopic connectivity* in a group of 17 participants at 1-month, 3-months, and 12-months after a single left-hemisphere stroke. These measures were also compared to a group of 13 age-matched healthy controls. The three measures exhibited different trajectories of change: (1) *modularity* steadily decreased across the 12-month period and became statistically inferior to control values at 12 months, indicating a less modular organization; (2) *hemispheric symmetry* values were abnormally low at 1-month and then increased significantly in the first 6 months, leveling off at levels not significantly below control levels by 12 months, suggesting that the two hemispheres diverged initially after the unilateral damage, but improved over time; and (3) *homotopic connectivity* exhibited a U-shaped function with a significant decrease from 1-6 months and then an increase from 6-12 months, to levels that were not significantly different from controls. The results revealed a complex picture of the dynamic changes the brain undergoes as it responds to abrupt onset damage.

## 1. Introduction

It is well understood that stroke in humans can affect neural network communication at multiple spatial scales. Micro changes impact small networks involving clusters of neurons while macro changes impact networks that support the communication between brain regions and communication across the hemispheres. In contrast to neuroimaging investigations that measure the impact of stroke in terms of changes in mean local activation levels within specific areas under specific task conditions, investigations of functional network properties examine the synchronization of activity across neuronal units. A variety of neuroimaging techniques (e.g., fMRI, MEG, EEG, electrocorticography) record the time-course of neural activity, although the most common approach to investigating functional connectivity uses resting-state fMRI (rsfMRI). rsfMRI provides a measure of the brain’s “intrinsic” functional connectivity that is not tied to a specific task. Correlations of the time-course of activities between brain areas, referred to as *functional connectivity (FC),* provide a basis for investigating the impact of stroke on functional network communication. Empirical and computational evidence clearly indicates that neural activities patterns are dynamic and that, when damaged, the human functional connectome undergoes neuroplastic changes over time (Klingbeil et al., 2019b; Sale et al., 2015; Saur et al., 2006; Siegel et al., 2018; Stam et al., 2010; Tao et al., 2022; Tao & Rapp, 2020, 2021). However, the effects of stroke on FC have been almost always examined at a single time-point subsequent to the stroke, and usually in the chronic stage (more than 1 year post stroke, see a review by Klingbeil et al. 2019). Thus, the nature, extent and trajectory of longitudinal FC changes subsequent to stroke have been scarcely examined. Moreover, most longitudinal studies evaluated treatment effects, comparing FC at pre and post-treatment time-points (e.g., Duncan & Small, 2016, 2018; Marangolo et al., 2016; Marcotte et al., 2013; Sandberg et al., 2015; Tao & Rapp, 2019, 2020; van Hees et al., 2014). Thus, there is a very significant gap in our understanding of the natural history of stroke on functional neural network communication.

In the work we report on here we examined three large-scale measures of functional connectivity: 1) *modularity*, a graph theoretic measure that quantifies the degree to which functional connections form modules of strongly interconnected (synchronized) brain regions (Rubinov & Sporns, 2010); 2) *hemispheric FC symmetry* that evaluates the degree to which the two hemisphere have similar patterns of intra-hemispheric FC; and 3) *homotopic FC* that characterizes the degree of communication (synchronization) between the hemispheres. These measures have been examined almost exclusively at the chronic stage of stroke (Duncan & Small, 2016; Tao et al., 2022; Tao & Rapp, 2019, 2020. But see Siegel et al. 2016 and 2018). Here we report on work that examined these three FC measures at three time-points during the first year after stroke (at approximately 1, 6, and 12 months post-stroke) in a longitudinal study in which stroke survivors participated in rsfMRI scanning at multiple time-points. This design allowed us to evaluate various possibilities regarding the trajectory of stroke-induced changes on FC, increasing our understanding of the degree to which brain injury impacts cerebral communication between and within hemispheres and how these change as recovery progresses. These understandings are not only important for developing theories of neuronal change after injury, but will inform interventional therapeutic approaches (e.g., when and where to apply brain stimulation)

### 1.1 Modularity: FC segregation into synchronized networks

The graph-theoretic measure of *modularity* quantifies the extent to which functional connections are segregated into tightly knit synchronized networks. Modularity is typically a single value computed over the whole brain or individual hemispheres. A widely used measure of modularity is Newman’s Q (Newman & Girvan, 2004) which quantifies the strength with which connections are grouped into networks relative to what would be expected by chance.

Evidence is mixed as to whether increased modularity is beneficial in both healthy and clinical populations. In healthy adults, modularity levels have been associated with, among other things, motor skill learning (Bassett et al., 2011, 2013); working memory (Braun et al., 2015; Kitzbichler et al., 2011; Yue et al., 2017), cognitive training (Gallen et al., 2016) and normal aging (Betzel et al., 2014; Chan et al., 2014; Meunier et al., 2009). Although higher modularity has often been often associated with better performance, one can imagine that higher modularity/system segregation might be more beneficial for certain tasks than others. In fact, Yue et al (2017) reported that, in healthy adults, higher modularity was associated with better performance on less complex tasks, while lower modularity (greater communication across different networks) was associated with better performance on more complex tasks. In a variety of neurological disorders, although higher modularity has been associated with less severe disease and stronger behavioral performance (Arnemann et al., 2015; Siegel et al., 2018; Tao & Rapp, 2019)), higher than normal levels of modularity have been reported in neurological disease (e.g., PD, MS; Baggio et al., 2014; Gamboa et al., 2014 respectively).

In the specific context of stroke, various issues have been investigated, including how modularity levels compare with those of healthy adults, the extent to which modularity is predictive of future responsiveness to treatment, and changes in modularity with treatment; very little work has examined the natural time-course of modularity changes subsequent to stroke. With regard to comparisons with controls both Siegel et al (2018) and Tao & Rapp (2019) reported comparable modularity levels in stroke and healthy controls, at the chronic stage. In terms of the relationship between modularity and recovery of function after stroke, the few studies that have examined the issue (Arnemann, et al., 2015; Duncan & Small, 2016; Siegel et al., 2018; Tao & Rapp, 2019) have all reported associations between higher pre-treatment modularity values and post-stroke behavioral improvement. For example, Arnemann et al (2015) found that pre-treatment modularity values predicted responsiveness to treatment (for attention and executive functions) with higher pre-treatment *modularity* predicting a better treatment response. This is consistent with findings in healthy adults in which baseline *modularity* values predicted responsiveness to behavioral intervention (Gall et al.,2015). Duncan and Small (2016) examined *modularity* and language performance before and after a period of language rehabilitation and found that they were positively correlated. Along similar lines, Tao and Rapp (2019) found that lower *modularity* levels prior to behavioral language therapy predicted a less robust response to treatment and, furthermore, they found that treatment resulted in increases in pre- to post-treatment *modularity* levels.

Specifically with regard to the longitudinal, natural time-course of *modularity* subsequent to stroke (the focus of the present study), to our knowledge only Siegel et al. (2018) evaluated this issue. They examined post-stroke *modularity* at 2 weeks, 3 months and 1 year after stroke, in a large sample (n=67 individuals with a range of lesion locations and deficit profiles, at the three timepoints). They found that at 2 weeks, modularity levels were reduced relative to healthy controls and then increased significantly after 2 weeks to almost normal levels by 3 months, with little change beyond that point. It is worth noting, however, that the *modularity* increases and normalization that Siegel et al. reported were observed only in the relatively small number of individuals who showed behavioral improvements over the same time period. In sum, the relatively small amount of available evidence highlights a complex picture of *modularity* changes following stroke and their relationship with behavior changes.

### 1.2. Left-right symmetry of hemispheric FC patterns

It has long been known that the two hemispheres are not structurally identical nor are they identical in terms of their instantiation of cognitive functions. In other words, there is well-established lateralization of function, such that (despite individual differences) certain functions tend to rely more on the left hemisphere (e.g., language) and others on the right (e.g., face recognition). It is assumed that the greater neural differentiation afforded by lateralization of function allows for specialization of function. Consistent with this, from birth to adulthood, brain areas supporting higher cognitive functions become increasingly lateralized (Dehaene-Lambertz et al., 2018; Olulade et al., 2020) as they become more specialized and efficient. However, even in healthy aging there is a point at which the tendency towards increasing lateralization is reversed (after late middle-age) and activation patterns then tend to become more distributed and less lateralized across the hemispheres (Cabeza, 2002; Meinzer et al., 2009), although it should be noted that the cognitive consequences of this decreasing lateralization are debated (Cabeza et al., 2018; Reuter-Lorenz & Park, 2010).

In terms of the brain’s response to neurological disease, different patterns have been reported for degenerative neurological disease compared to stroke. A reduction in hemispheric specificity compared to healthy controls and, thus a greater “homogenization” of activity patterns in the hemispheres has been reported in primary progressive aphasia (Tao et al., 2022). For stroke, in contrast, there have been a number of reports that the ipsi-lesional and contra-lesional hemispheres undergo different activation and FC changes between and within specific networks and regions, revealing considerable post-stroke asymmetry in hemispheric FC networks (Hartwigsen & Saur, 2019; Sandberg, 2017; Saur et al., 2006). In fact, Tao et al (2022) carried out a direct comparison of FC characteristics in PPA and post-stroke groups matched for language profiles and lesion epicenter (posterior frontal, anterior parietal and insula). For each group, they compared the two hemispheres on a number of intra-hemispheric FC measures and found that, while the neurodegenerative PPA group showed increased similarity of the two hemisphere’s FC patterns relative to healthy controls, for the stroke group the FC patterns of the two hemispheres were less similar to each other than they were in controls.

With regard to the longitudinal consequences of stroke, the available data suggest a reduction in the FC symmetry of the hemispheres relative to controls and the asymmetry may lessen over time (e.g., Saur et al, 2006). However, very few studies (except Tao et al. 2022) have directly examined this issue, and thus the progression of FC symmetry changes in response to stroke during recovery is not understood.

### 1.3. Synchronization of neural activity between the hemispheres: Homotopic FC

Distinct from the question of the similarity of the FC patterns within the two hemispheres is the question of the degree to which the hemispheres communicate with one another. Inter-hemispheric communication (the majority of which takes place between homotopic areas) has been shown to play an important role in a range of cognitive processes (Gotts et al., 2013; Gracia-Tabuenca et al., 2018; Jin et al., 2020; Zhao et al., 2020). The structural basis for interhemispheric communication is largely the corpus callosum, whose fibers primarily connect homologous regions of the two hemispheres (Eccer, 2014).)

Homotopic FC specifically measures the degree of correlation (synchronization) of the time-series of BOLD responses between pairs of homotopic brain regions using data from either rsfMRI and or background FC (from task-based fMRI, Cole et al., 2014). The reliance of homotopic FC on the corpus callosum is supported by findings that corpus callosum lesions are associated with decreased magnitude of homotopic FC (Gracia-Tabuenca et al., 2018; Tzourio-Mazoyer et al., 2016). Homotopic FC has been shown to index functions such as attention, executive functions, and memory (Chen et al., 2019). Studies of homotopic FC in neurotypical individuals over the lifespan (e.g., (Tzourio-Mazoyer et al., 2016; Zhao et al., 2020; Zuo et al., 2010) reveal various trajectories of longitudinal change in homotopic FC that are influenced not only by age, but also gender and brain region.

Specifically with regard to the effects of stroke on inter-hemispheric communication, Siegel et al (2016) and Tao & Rapp (2020) found that lower levels of homotopic FC were associated with greater deficit severity in multiple cognitive domains, consistent with previous findings in the motor and attention domains (Baldassarre et al., 2016; He et al., 2007; Rehme et al., 2015; van Meer et al., 2012). In terms of comparisons with controls, Siegel et al. (2016) evaluated homotopic rsfMRI sub-acutely (2 weeks) after stroke (n=132; left and right hemisphere stroke). On the basis of their findings, they proposed that reduced synchronicity of functional brain responses between the hemispheres relative to controls should be considered a “general physiological network phenotype of stroke”. In their review of FC studies in post-stroke aphasia, Klingbeil et al (2019) found general support for the Siegel, et al. claim (see also, New et al., 2015; Sandburg, 2017; Zhu et al., 2014). The Siegel et al. claim was subsequently supported by the Tao and Rapp (2020, 2022) findings of reduced homotopic FC relative to controls at a very chronic post-stroke stage (> 6 month time post stroke).

In terms of the natural time-course of homotopic FC changes, as opposed to treatment-induced changes, subsequent to stroke, to date there are nolo ngitudinal studies. Thus, while reduced homotopic FC has been reported at both sub-acute and chronic stages after stroke, the time-course between these time-points has not been described, leaving open various possibilities, including increasing, decreasing or nonlinear trajectories of homotopic FC change over this time period, reflecting potential windows of neuroplasticity during recovery.

### 1.4 Current Study

A review of the literature reveals that the consequences of stroke for large-scale functional connectivity properties have been examined almost entirely at the chronic state and even then, there have been very few studies. The picture that emerges is one of reduction, in comparison to healthy controls, in both the synchronicity of homotopic communication and the symmetry of intra-hemispheric FC patterns across the two hemispheres. Modularity has also been found to be reduced immediately following stroke and then followed by recovery of normal levels, at least under some conditions.

The current study aimed to address the gaps, described above, in our understanding of the functional connectivity changes that occur in the first year subsequent to a unilateral left hemisphere stroke. Using longitudinal resting-state fMRI data collected in a group of individuals with unilateral left-hemisphere stroke at approximately 1, 6 months, and 12 months post-stroke, we examined the temporal trajectories of three large-scale FC properties: *modularity*, *hemispheric FC symmetry* and *homotopic FC*. The analyses reveal significant changes in all three FC network characteristics during the first post-stroke year with distinctive trajectories for each. The findings underscore the dynamic nature of the brain’s response to sudden onset damage and provide a foundation for future investigations of the consequences of stroke for the brain’s functional network organization.

## 2. Methods

### 2.1 Participants

This retrospective study was part of a larger ongoing project involving multiple comprehensive stroke centers in Houston, TX, USA that consecutively recruited monolingual English speakers with left hemisphere acute stroke independently of aphasia diagnosis without other health conditions that could impact cognition (i.e., tumor, dementia, alcohol and/or drug dependency). Twenty-one of the recruited participants returned for neuroimaging for at least two of three timepoints during the year after stroke (1 month, 6 months, 1-year post-stroke). In the returning sample, we further excluded three participants with brain stem stroke and one participant with cerebellar stroke, resulting in a total of 17 participants in the current study (Age: M=59, SD=8.48, range 46-73 years; Education: M=15, SD=3.07, range 12-23 years; 11 females; 3 left-handed; 8 received clinical diagnosis of aphasia). Among the 17 participants, 10 had imaging data available from the first two time points (1 and 6 months), 14 from the 2^nd^ and 3^rd^ timepoints (6 and 12 months), and 7 from the three time-points (see Table 1). Detailed information about the participants is reported in Table 1. We also collected imaging data from 13 healthy controls (mean age=55.5, SD=14.2 years, mean education=15.9, SD=2.6 years, 10 Females, all but 2 were right-handed).

**Table 1.** Demographic information.

| **Participant** | **Age** | **Education** | **Gender** | **Right-handed** | **Lesion volume (cm<sup>3</sup>)** | **Scanning time-points** |
| --- | --- | --- | --- | --- | --- | --- |
| **1** | 65 | 12 | F | Yes | 3.56 | 1mo, 6mo |
| **2** | 47 | 12 | F | Yes | 3.32 | 1mo, 6mo |
| **3** | 70 | 14 | F | Yes | 2.84 | 1mo, 6mo |
| **4** | 73 | 16 | M | Yes | 8.05 | 6mo, 12mo |
| **5** | 63 | 18 | M | Yes | 4.94 | 6mo, 12mo |
| **6** | 63 | 23 | F | Yes | 11.58 | 6mo, 12mo |
| **7** | 53 | n/a | M | No | 3.75 | 6mo, 12mo |
| **8** | 49 | 14 | F | No | 0.11 | 6mo, 12mo |
| **9** | 46 | 16 | M | Yes | 5.27 | 6mo, 12mo |
| **10** | 56 | 16 | F | Yes | 0.22 | 6mo, 12mo |
| **11** | 59 | 12 | M | Yes | 9.13 | 1mo, 6mo, 12mo |
| **12** | 53 | 12 | F | Yes | 10.31 | 1mo, 6mo, 12mo |
| **13** | 61 | 15 | F | Yes | 22.38 | 1mo, 6mo, 12mo |
| **14** | 59 | 12 | F | No | 12.26 | 1mo, 6mo, 12mo |
| **15** | 70 | 16 | F | Yes | 2.59 | 1mo, 6mo, 12mo |
| **16** | 66 | 18 | F | Yes | 2.46 | 1mo, 6mo, 12mo |
| **17** | 73 | 12 | M | Yes | 3.08 | 1mo, 6mo, 12mo |

Due to the characteristics of the larger project, behavioral assessment was limited to language tasks and, thus, behavioral measures of general cognitive abilities were not collected. As a result, we did not have the behavioral measures appropriate for analyses examining the relationship between global measures of functional connectivity and behavior. We discuss this limitation in the General Discussion.

Lesion masks were manually drawn on T1 images using ITK-snap (Yushkevich et al., 2006). We registered T1 images to the Colin-27 template using ANTs and normalized masks to the MNI space based on the transformation parameters reported in Avants et al. (2008) (see Ding & Schnur, 2022 for a similar approach). We created one lesion mask for each participant by taking the union of their lesion tracing masks across scans obtained at all available time-points. These masks were used for subsequent analyses. As seen in Fig. 1, the participants exhibited diverse lesion distributions (the maximum number of participants with lesions at a given voxel was four (23.5% of the cohort), ranging from restricted white matter or subcortical lesions to typical MCA stroke lesion patterns. Across the 17 participants, the lesions had average volumes of 6.23 cubic centimeters (SD=5.43, range 0.11 – 22.38).

**Figure 1.**
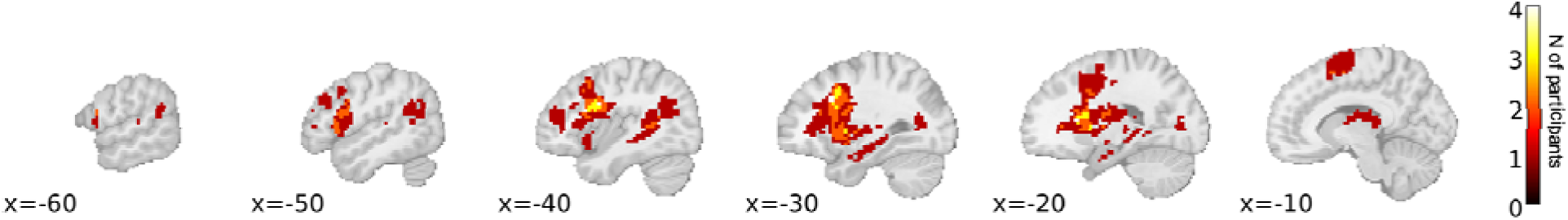
Lesion overlap. The color scale indicates the number of participants in whom the voxel was lesioned.

### 2.2. Imaging acquisition

Due to an extended participant recruitment period, two different sets of T1 and rs-fMRI scanning sequences were used in a Philips or Siemens PRISMA 3T scanner (see below). Importantly, however, for each participant, the scans at different time-points used the same sequence parameters. The two sequences are as follows. Sequence A: Phillips Ingenia 3T; rs-fMRI: TR/TE=2000/30ms, Flip angle=90 deg, voxel size=2.75*2.75*3mm, FOV=220mm, volume number=150; T1: TR/TE=8.4/3.9ms, Flip angle=8 deg, voxel size=1*1*1mm, FOV=240mm, slice number=175. Sequence B: Siemen’s Prisma 3T; rs-fMRI: TR/TE=1200/30ms, flip angle=90 deg, voxel size=2*2*2mm, FOV=192mm, volume number=500; T1: TR/TE=2600/3ms, flip angle=8 deg, voxel size=1*1*1mm, FOV=256mm, slice number=176. In total, Sequence A was used with four participants and Sequence B with 13 participants.

### 2.3. Imaging data preprocessing

We used DPABI V4.2 to automatically preprocess the rs-fMRI data (Yan et al., 2016) We removed the first 10 time-points of data, corrected slice-timing, realigned all the volumes, regressed out the average signals of the white matter, the CSF, and 24 head-motion parameters, normalized functional images using the transformation information derived from the segmentation of co-registered T1 images, smoothed data with a 4 FWHM Gaussian kernel, and filtered frequency signals (0.01-0.1 Hz).

### 2.4. Estimating functional connectivity (FC)

Functional connectivity matrices were constructed using an atlas consisting of 200 pairs of homotopic cortical gray matter parcels (Yan et al., 2023). We used this atlas because a key analysis specifically concerned the relationship between the two hemispheres. FC values were estimated as Pearson correlations (with Fisher’s transform) of the timeseries. For network-based measures (system segregation and modularity), we used the 7-network organization provided with the atlas (Yeo et al., 2011; Yan et al., 2023). In this atlas, the 200 parcels were organized into seven functional networks (see Fig. 2a): dorsal attention network (DAN), sensorimotor network (SMN), visual, ventral attention network (VAN), default-mode network (DMN), limbic, and frontoparietal network (FPN). Four of the networks, DAN, VAN, DMN, and FPN are distributed across the frontal, parietal, and temporal lobes, SMN and Visual networks are mainly located in sensorimotor and

**Figure 2.**
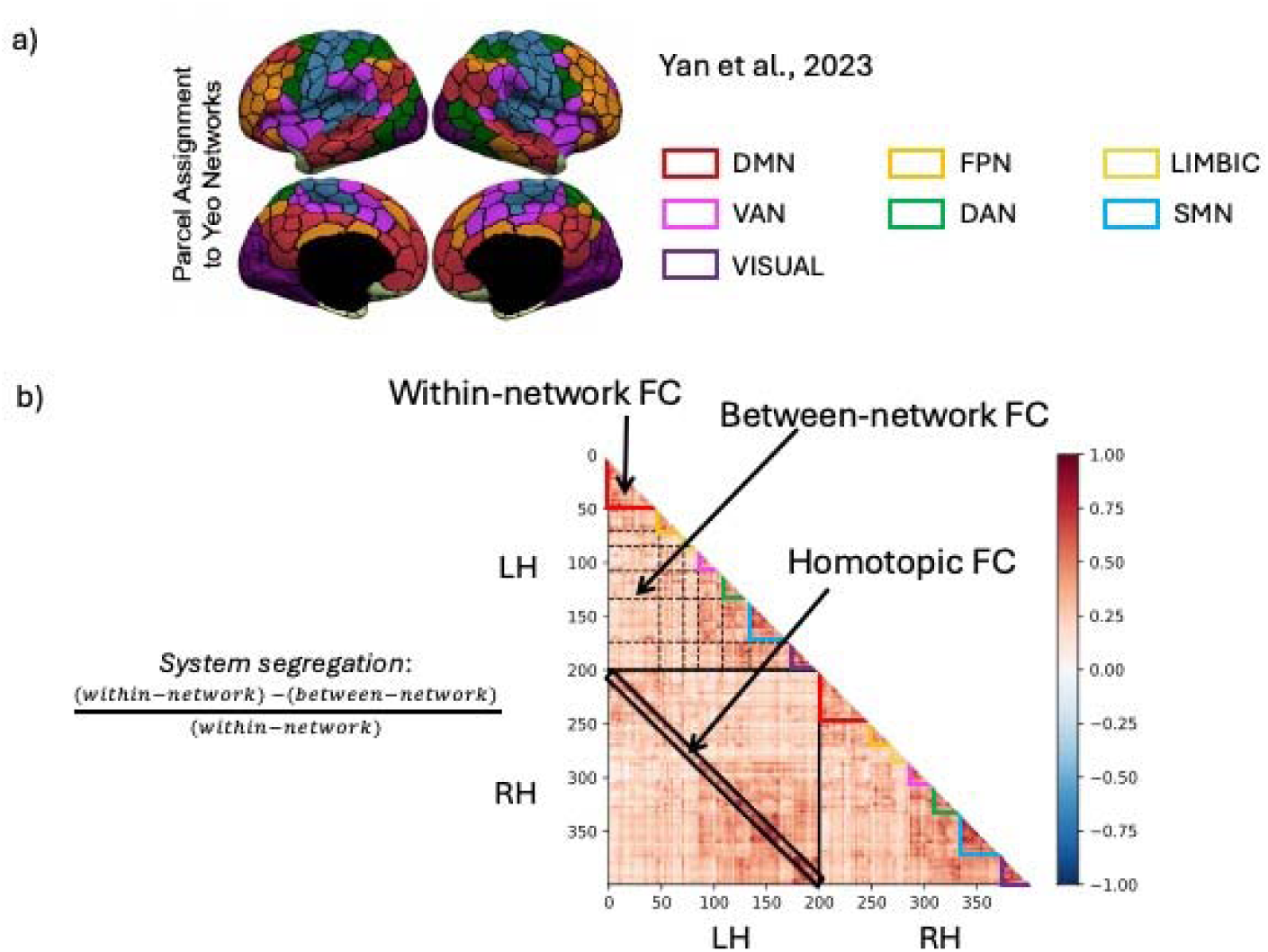
The resting-state networks used in the analyses and a sample functional connectivity matrix. (a) The (color-coded) parcels that make up the seven resting-state networks defined by Yan et al. (2023) and used in subsequent analyses; (b) A pair-wise connectivity matrix is shown that contains the functional connectivity (FC) of the 400 parcels (200 in each hemisphere) from a homotopic atlas proposed by Yan et al. (2023). Note that the full matrix is fully diagonally symmetrical so we display only half of the matrix. The lower left quadrant contains the inter-hemispheric FC values with the *homotopic FC* values indicated within the black outlined diagonal (i.e., connections between pairs of homotopic parcels). The upper left and lower right quadrants contain intra-hemispheric FC values. The intra-hemispheric connections are divided into within- and between-network functional connections based on the pre-defined seven networks shown in (a) *System segregation* was quantified as a ratio between the within- and between-network FC, following Chan et al. (2014). *DMN: Default-mode network; VAN: ventral attention network; FPN: Frontoparietal network; DAN: Dorsal attention network; SMN: Sensorimotor network*.

Occipital cortices, respectively. and the Limbic network included the temporal poles and orbital frontal areas.

### 2.5. Functional connectivity (FC) measures

As described in the Introduction, we computed three FC measures that were used in the subsequent analyses. These were computed for each participant, at each time-point at which they were scanned (1, 6 and/or 12 months post stroke): (1) modularity, (2) hemispheric FC symmetry and (3) homotopic FC.

(1) *modularity* is a graph-theoretic measure that quantifies the magnitude of the pairwise connectivity strength of nodes within the 7 networks (see Fig. 2a) relative to what would be expected by chance given the total number of connections (Rubinov and Sporn, 2008). Here we specifically computed weighted *modularity* with the Python toolbox *bctpy* (https://pypi.org/project/bctpy/). In addition to whole brain modularity, we also calculated *modularity* within each hemisphere using only intra-hemispheric connections to evaluate possible hemispheric differences^1^.
(2) For *hemispheric symmetry* we measured the similarity of FC patterns across the 200 parcels in each of the two hemispheres. We specifically examined symmetry in terms of the measure of *system segregation* which is computed, for each parcel, as the ratio of the within- and between-network FC (see Fig. 2b), with “network” referring to the 7 networks listed above. Then, for each participant, *hemispheric symmetry* was quantified as the Pearson correlation between the system segregation values of the 200 parcels in each hemisphere, yielding one *hemispheric symmetry* value per participant at each time-point.
(3) For *homotopic FC*, for each participant we calculated *homotopic FC* as the connectivity strength between the time-courses of each of the 200 pairs of homotopic parcels, resulting in 200 values per participant (Fig. 2b).

### 2.6. Statistical analyses

For each FC measure, we evaluated longitudinal changes across the three time-points using separate linear mixed effects models for each of the three FC measures. For *modularity* and *hemispheric symmetry*, the dependent measure consisted of the FC measures for healthy controls (one time-point only) and the stroke participants (for each time-point at which they were scanned). *Time-Point* was the fixed effect of main interest. Controls were coded as 0 while stroke participant data was coded for time-point (1, 2, and/or 3) so that stroke participant values at each time-point could be directly compared to the control values. *In-scanner motion* (average RMS) was included as the covariate and *Participant* served as the random effect. For *homotopic* FC, the same LMEM structure was used except that the connectivity strengths of each of the 200 homotopic connections for each participant and time-point were all entered into the model, and an additional random effect of *Connection* for each of the 200 connections was included.

We examined longitudinal changes by contrasting each pair of time-points: 1-month vs. 6-months, 6-months vs. 12-months, 1-month vs. 12-months, and also compared the stroke participants’ values of each time-point to the values obtained from Control participants to evaluate the magnitude of the stroke participant effects as well as the directionality of the longitudinal trajectories relative to normal levels.

Because we had a mixed longitudinal/cross-sectional design (not all participants were scanned at the same time-points), to evaluate if the effects for the full participant group could have been carried by the subset of participants who only had data for two time points, we repeated all the above analyses for the participants who had been scanned at all the three-timepoints (N=7). Moving forward, we refer to this fully longitudinal group as the “sub-group” and for each of the analyses, we first report results from the whole group and then for the sub-group.

We also examined, in a separate set of analyses, if lesion size was related to any of the FC measures. To do so, we used the same regression models as described above except that 1) healthy controls were not included, 2) lesion size (log transformed) was included as a predictor, and 3) the interaction of lesion size by time-point was included. None of the analyses showed significant effects of lesion size and these specific results are reported in Supplementary Materials.

## 3. Results

### 3.1. Longitudinal changes in *modularity*

As depicted in Figure 3, whole-brain m*odularity* showed a significant overall decrease from 1- to 12-months (t=-2.66, p=0.013, Fig. 3a) with similar but non-significant decreases in both sub-periods (1- to 6-month: t=-1.33, p=0.197, 6- to 12-month: t=-1.71, p=0.102). Thus, overall, networks became increasingly less autonomous and more interactive. When compared to Controls, *modularity* values were numerically lower over the first year, although only the values at the 12-month assessment were significantly lower than controls (1-months vs. controls: t=-0.21, p=0.835; 6-months vs controls: t=-1.13, p=0.268; 12-months vs. controls: t=-2.15, p=0.038, Fig. 3a).

**Figure 3.**
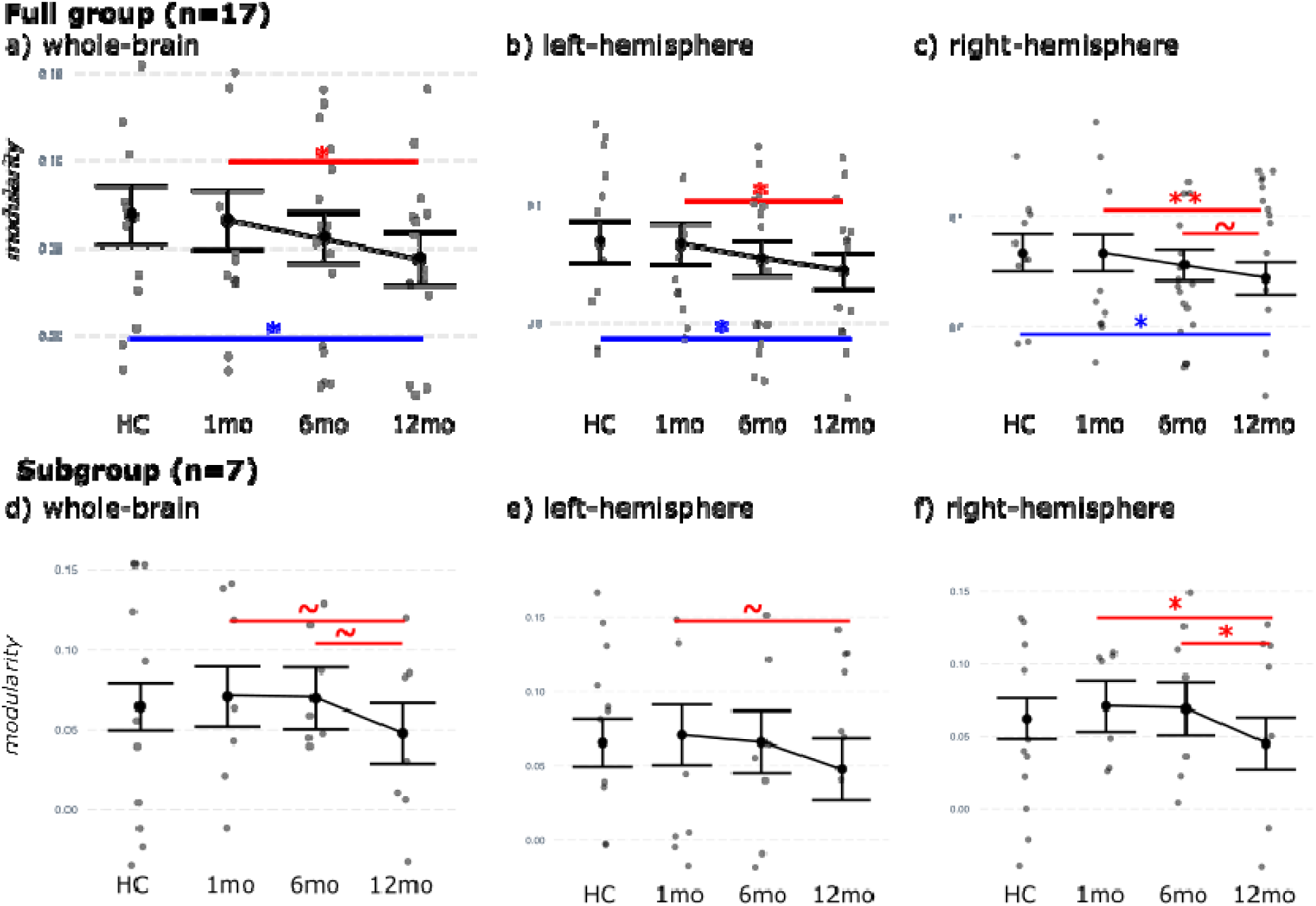
Longitudinal changes in *modularity*. a)-c) Results from the whole group (n=17), d)-f) Results from the subgroup of 7 individuals scanned at all three time-points. Left to right: *modularity* calculated over the whole-brain and within each hemisphere. Statistically significant longitudinal effects for time-point comparisons indicated with red lines whose end points mark the relevant time points. Statistically significant effects for comparisons of the stroke and healthy control groups (HC) are marked in blue. *\*\*:p<0.01, *:p<0.05, ∼:p<0.1*

For each of the hemispheres, *modularity* values showed patterns of longitudinal decrease that were similar to what was observed for the whole brain (Fig. 3b and 3c) (Left: 1- to 12-months: t=-2.60, p=0.015; 1- to 6-months: t=-1.52, p=0.141; 6- to 12-months: t=-1.40, p=0.176;. Right: 1- to 12-months: t=-2.88, p=0.008; 1- to 6-months: t=-1.42, p=0.169; 6- to 12-month: t=-1.88, p=0.073). Compared to controls, the patterns were again similar as in the whole brain, with patient participant values significantly lower than those of the controls at 12-months (Left: 1-month vs. controls: t=-0.15, p=0.884; 6-months vs controls: t=-1.22, p=0.232; 12-months vs. controls: t=-2.07, p=0.045. Right: 1-months vs. controls: t=-0.01, p=0.99; 6-months vs controls: t=-0.95, p=0.348; 12-monthsvs. controls: t=-2.04, p=0.049).

These longitudinal pattern of decreasing *modularity* were confirmed in the subgroup analysis (Fig. 3d-f, 1- to 12-months: t=-2.11, p=0.054; 1- to 6-months: t=-0.10, p=0.926; 6- to 12-months: t=-2.01, p=0.064). Compared to controls, values for the subgroup were numerically but not statistically different (1-month vs. controls: t=0.53, p=0.604; 6-months vs controls: t=0.42, p=0.679; 12-months vs. controls: t=-1.38, p=0.181, Fig. 3d).

For this subgroup, the hemispheric effects also showed the same pattern as the whole-brain effects (See Fig. 3e, 3f and Table S1).

Overall, the findings indicate that, in the first year following a left-hemisphere stroke, the brain’s modular organization increasingly deviated from that of the controls with **functional modules becoming increasingly less segregated from one another and, thus, more interactive.**

### 3.2. Longitudinal changes in *hemispheric symmetry*

*Hemispheric symmetry* quantifies the degree of similarity between the two hemispheres in terms of hemisphere-internal FC patterns. We specifically evaluated symmetry in terms of the distribution of the FC measure of *system segregation* (Chan et al., 2014, see Fig. 2b) across the parcels of each hemisphere. This symmetry value, provides a measure of FC organization on a continuum from highly lateralized (asymmetrical) to fully bilateral (symmetrical).

The distributions of *system segregation* values for all of the parcels in the two hemispheres across time-points are visualized in Figure 4a. The results of the statistical analyses for the whole group are depicted in Figure 4b. For this group, *Hemispheric symmetry* increased significantly from 1- to 6-months (t=2.41, p=0.022) and then values remained relatively stable from 6- to 12-month (t=-0.47, p=0.645); 12-month values were marginally higher than those at 1-month (t=1.90, p=0.066). In addition, when compared to the controls, patient participant values at 1-month were significantly lower, but at 6- and 12-months FC symmetry was not different from what was observed in the controls (1-month vs. controls: t=-2.24, p=0.030; 6-month vs controls: t=-0.44, p=0.666; 12-month vs. controls: t=-0.77, p=0.444, Fig. 4b).

**Figure 4.**
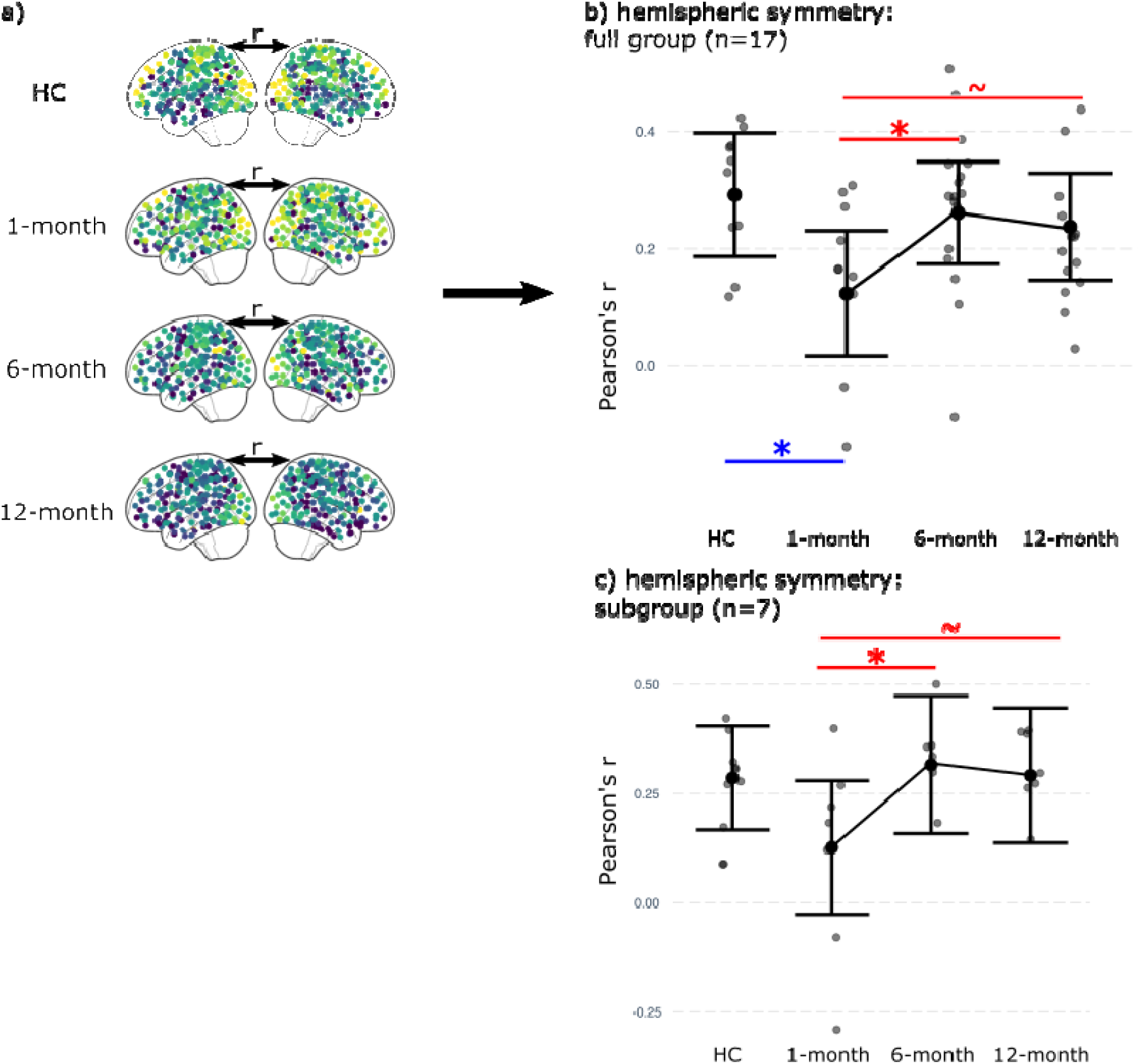
Longitudinal changes in *hemispheric symmetry* of the FC measure of *system segregation.* (a) The distributions of parcel-wise values of *system segregation* averaged across the healthy controls (HC) and stroke participants at each time-point. *Hemispheric symmetry* was quantified as the Pearson correlation (r) of the *system segregation* values between the two hemispheres; (b) Results of the statistical analysis of *hemispheric symmetry* for the whole group; (c) Results for the subgroup of 7 individuals scanned at all three timepoints. Statistically significant longitudinal effects for time-point comparisons are indicated with red lines whose end points mark the relevant time points, while results of statistically significant effects for comparisons of the stroke and healthy control groups (HC) are indicated in blue. *\*: p<0.05, **:p<0.01, ∼:p<0.1*

The effects were similar in the subgroup data (Fig. 4c) where hemispheric symmetry increased between 1 and 12-months (t=2.11, p=0.053); this included a significant increase between 1 and 6-months (t=2.41, p=0.030), and then values remained relatively stable from 6- to 12-months: t=-0.31, p=0.759).). The comparisons to the control group showed the same numeric pattern, although the effects were not statistically significant (1-month vs. controls: t=-1.65, p=0.114; 6-months vs controls: t=0.30, p=0.767; 12-months vs. controls: t=0.06, p=0.953, Fig. 4c).

Overall, the data show that *hemispheric symmetry*, the similarity between the hemispheres with regard to the distribution of *system segregation* within each hemisphere, was significantly reduced **early after stroke, such that the two hemispheres exhibited less similar organization patterns than observed in controls, and that these patterns then approached normal levels of symmetry (lateralization) after about 6-months.**

### 3.3. Longitudinal changes in homotopic FC

*Homotopic FC* evaluates the extent of FC synchronization between the hemispheres. As can be seen in Figure 5, *homotopic FC* values showed a U-shaped response such that *homotopic FC* significantly decreased from 1 to 6-months post-stroke (t=-2.81, p=0.005) and then significantly increased from 6-months to 12-months (t=7.09, p=1.39e-12), with values at 12-months significantly higher than at the 1-month timepoint (t=2.95, p=0.003). When compared to controls, all three time-points were numerically lower, although not statistically so (1-month vs. controls: t=-1.22, p=0.234; 6-months vs controls: t=-1.54, p=0.135; 12-months vs. controls: t=-0.86, p=0.399).

**Figure 5.**
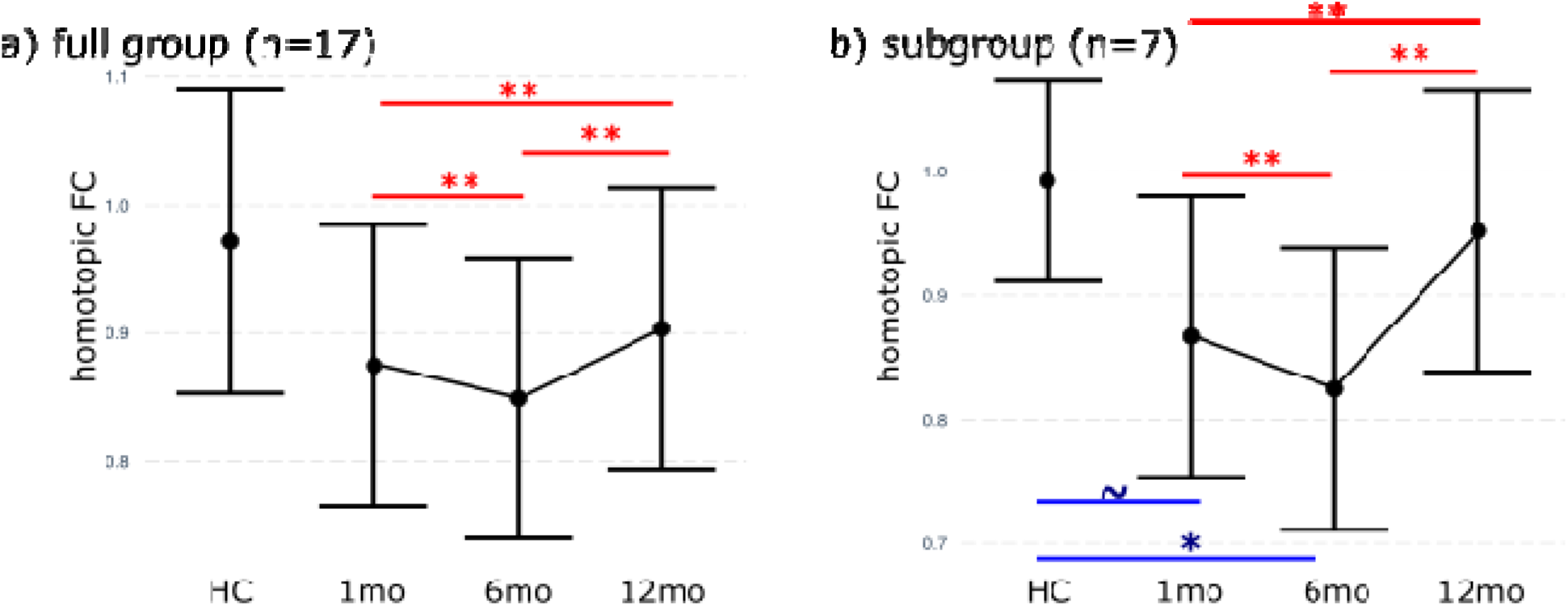
Longitudinal changes in *homotopic functional connectivity* (FC) in the first year post-stroke. a) Results of the statistical analyses for the entire group and b) Results of the statistical analyses for the subgroup group of 7 stroke participants who were scanned at all three time-points. For both groups, *homotopic FC* first decreased from 1- to 6-months, then increased from 6- to 12-months, approaching normal levels. Statistically significant effects for time-point comparisons are indicated with red lines whose end points mark the relevant time points, while results of statistically significant effects for comparisons of the stroke and healthy control groups (HC) are marked in blue. *\*: p<0.05, **:p<0.01, ∼:p<0.1*

These patterns were confirmed by the analysis of the subgroup. Like the larger group, homotopic FC significantly decreased from 1 to 6-months post-stroke (t=3.75, p=0.0002) and then significantly increased from 6-months to 12-months (t=11.235, p<2e-16), with the values at 12-months significantly higher than at the 1-month timepoint (t=7.47, p=9.7e-14). In the subgroup, the homotopic FC values were significantly lower than those of controls at the first two time-points (1-month vs. controls: t=-1.87, p=0.081; 6-months vs controls: t=-2.50, p=0.024; 12-months vs. controls: t=-0.60, p=0.555).

Overall, the longitudinal patterns indicate that **the two hemispheres became less synchronized with each other in terms of interhemispheric, homotopic FC in the first six months after stroke, and then approached normal levels.**

## 4. General Discussion

We report on an investigation of the longitudinal time-course of functional network connectivity changes in the first year following a left hemisphere stroke. Seventeen stroke survivors each participated in two or three fMRI scanning assessments at approximately 1, 6 or 12 months subsequent to a stroke, allowing an examination of changes from the sub-acute to early chronic stage. Thirteen control participants were also scanned at a single time-point. From resting-state functional MRI (rsfMRI) data we calculated functional connectivity (FC) measures to compare across time points for the stroke participants as well as between stroke and control groups. FC connectivity (the degree of synchronization of the BOLD time-course) between brain areas has been shown, in a large number of studies, to be associated with various behaviors and measures of recovery of function in the context of stroke (see review by Klingbeil et al., 2019a). While there are a large number of FC measures that can be computed, here we specifically analyzed three large-scale, whole-brain measures: *modularity*, *hemispheric symmetry* (of intra-hemispheric *system segregation* values) and *homotopic FC* (synchronization of homotopic regions). Prior research examining the post-stroke longitudinal changes of FC network properties is scant and largely focused on specific brain regions (Nair et al., 2015; Sebastian et al., 2016; Zhu et al., 2014). With regard to longitudinal changes, to our knowledge, only Siegel et al. (2018) carried out a longitudinal examination of *modularity* in the first year subsequent to stroke. The longitudinal effects of stroke on hemispheric symmetry and homotopic FC are largely unknown. The current study aimed to address these gaps and provide a foundation for future investigation.

The data analyses revealed different trajectories of longitudinal FC change for the three measures, underscoring the complexity of the neurodynamic consequences of stroke. The following are the key findings: (1) the *modality* of functional networks steadily decreased across the 12-month period, such that *modularity* values at 12 months were inferior to those of controls; (2) *hemispheric symmetry* values were abnormally low at 1 month and then increased significantly in the first 6 months, leveling off at levels not significantly below that of controls by 12 months; and (3) *homotopic connectivity* values exhibited a u-shaped function with a significant decrease from 1-6 months and then an increase from 6-12 months, to levels that were not significantly different from controls. Comparable patterns were found for the subgroup of participants that all participated in three scanning sessions. In the following sections we discuss how these findings compare to those reported in the literature, finishing with a discussion of the limitations of this study.

### 4.1. Relationship to previous literature

#### Modularity

The finding of steadily decreasing and lower than normal levels of *modularity* at 12 months is at odds with the findings of Siegel et al. (2018) and Tao & Rapp (2019) who found *modularity* levels comparable to controls in the chronic post-stroke stage. Tao & Rapp’s participants were mostly quite chronic (mean=58 months, range 14-119), leaving open the possibility that normal levels are achieved only much later. Siegel et al. found that while *modularity* was below normal acutely (at 2 weeks post stroke) it returned to almost normal levels by 3 months and showed little change beyond that point. However, Siegel et al. (2018) reported this pattern of normalization was driven by the subgroups of individuals (n= 5-23 depending on the task) who showed relatively good behavioral recovery in cognitive tasks (attention, spatial, verbal memory and language). For the group that they considered to show “no recovery” (n=11-22) in behavior over the year, *modularity* changes were at best extremely modest. This leaves open the possibility that, indeed, modularity normalization may typically take place over a long period of time, except in those individuals who exhibit rapid behavioral improvement. This relationship with modularity and behavioral performance increases would be consistent with the finding that positive treatment effects are most often associated with increases in modularity (Tao & Rapp, 2019; Duncan & Small, 2016) and more generally, in healthy participants, that higher modularity values are associated with better learning outcomes (Bassett et al., 2011, 2013; Kitzbichler et al., 2011 and others).

However, as noted in the Introduction, task complexity may determine the “ideal” amount of system *modularity,* such that more complex tasks may benefit from lower modularity -less system segregation and more interaction between modules (Yue et al., 2017). Given that finding, it is possible that decreasing modularity over the first year (for at least many stroke survivors) reflects the complexity of the cognitive demands they encounter in the face of the cognitive disruptions caused by the stroke. In that case, decreasing modularity could actually be adaptive as the brain attempts to increase communication between modules in an effort to solve the challenges it faces. Perhaps, only with time and the rebuilding/restructuring of the functional modules do normal levels of segregation and specialization return. Under that account, we would expect normal levels of *modularity* to return at different points in time for different individuals, in line with their pace of functional recovery.

#### Hemispheric FC symmetry

In the current study, we quantified the FC symmetry of the two hemispheres by considering the measure of *system segregation* (the ratio of average within network/between network FC strength for each region; Chan et al., 2014, see Figure 2b). We chose this measure because it represents global summary of a region’s FC characteristic. Hemispheric symmetry was measured as how similar *system segregation* patterns were in the two hemispheres. This provides a measure of the extent of FC lateralization, characterizing the general degree of internal similarity between the hemispheres. While, as reviewed in the Introduction, several studies have reported differences in post-stroke FC characteristics of the two hemispheres (e.g., Hartwigsen & Saur, 2019), to our knowledge it is only Tao et al. (2022) who carried out a direct comparison of global hemispheric FC organization, allowing an assessment of FC symmetry. Tao et al. (2022) evaluated four different measures, similar to the *system segregation* measure considered here, and found that in chronic stroke (mean of 6.6 years post stroke, SD = 4.7), for all measures, FC symmetry was significantly reduced relative to controls. In other words, the hemispheres differed from each other to a greater extent after stroke (consistent with more functional lateralization), at least in the very chronic stage. This contrasts with the results of the current study, in which although we also found decreased FC symmetry in the stroke group, it was statistically significant only at 1 month post stroke and then increased and was numerically, but not statistically, inferior to the control group by 6 months, a level which was maintained at 12 months post stroke.

In terms of the adaptiveness of these changes with regard to behavior, as reviewed in the Introduction, it has been found that decreasing hemispheric symmetry is associated with normal healthy development and specialization of function, while with aging and neurodegenerative disease increases in symmetry (homogenization) are observed. This suggests that asymmetry may be positive at least under certain circumstances and increased symmetry may be associated with cognitive decline. However, a specific examination of the relationship between abnormally high levels of hemispheric asymmetry (as observed in stroke) and behavior has not been reported, and we were not able to evaluate this important question ourselves in this study.

Another question raised by this study is: How can we account for the apparent “normalization” of FC symmetry by 6 months post-stroke in the current study while Tao et al., (2022) did not observe this even in the highly chronic stage? We do not have sufficient information to know to what extent the differences in FC symmetry findings between these studies are related to differences between the samples in terms of neural or other factors. In future work it will be especially important to consider the relevance of lesion location give that this varied across the left hemisphere in the current study, while (by design) lesions were concentrated in the left IFG and insula in Tao et al (2022).

#### Homotopic functional connectivity

As discussed in the Introduction, Siegel et al (2016) proposed that reduced homotopic synchronicity should be considered a “general physiological network phenotype of stroke”, which they observed at 2 weeks post stroke in a large sample of left and right hemisphere stroke patients (n=100). This general claim was subsequently supported by Tao and colleagues (2020, 2022, see also Klingbeil et al., 2019) who also reported reduction in homotopic FC, although, in contrast to Siegel 2016, 21018) they examined a chronic sample at several years post stroke. In the current study, we found that homotopic FC values were reduced relative to controls at 1, 6, and 12 months post-stroke, although these effects were statistically significant only at 1 and 6 months. Thus, on the one hand, we confirm the overall finding of reduced homotopic FC in stroke, on the other hand we find numerically but not statistically significant effects at the chronic stage. The latter, however, could be a consequence of low power and, thus, confirmation awaits future evaluation with more highly powered studies.

The U-shaped time-course of homotopic FC connectivity in the current sample was noteworthy in showing an initial significant decline and followed by a significant increase (towards normalization). The decrease in homotopic connectivity from 1 to 6 months is particularly intriguing. It suggests lowered homotopic FC may not only be an immediate response to stroke (as reported by Siegel et al., 2016), but may be an active process, potentially associated with the brain’s attempt to optimally respond to the neural insult and reduction in representational and processing capacity. Under those circumstances, further isolating the hemispheres could be an important first step in which hemisphere-internal reorganization takes place, subsequently followed by re-establishing more normal levels of inter-hemispheric communication.

Overall, what is clear from the findings of this study is that these three FC measures a) are highly dynamic in the first year post-stroke and b) exhibit quite different trajectories of change during that year. While both hemispheric symmetry and homotopic communication move sharply from depressed levels to normalization, they do so at different timepoints, with the inflection point for *hemispheric symmetry* occurring at 1 month and for *homotopic connectivity* at 6 months. Further, in our sample, we don’t see a “normalization inflection point” for *modularity;* presumably it may occur at an even later timepoint. These findings raise a number of questions, among others: Is this the necessary order of normalization? Are these effects sequentially or otherwise dependent on one another? Is there a unifying principle/s that accounts for these time-courses? These are all critical questions to be answered with larger samples and presumably with sophisticated computer modeling to aid in hypothesis testing.

### 4.2 Limitations

This study has several limitations. First among them is the relatively small sample size. It is challenging to carry out longitudinal investigations beginning at the sub-acute stage after stroke, involving multiple scanning sessions over an extended period. For this reason, there have been so few (to our knowledge only Seigel et al., 2016) longitudinal studies of FC in the first year after stroke. Nonetheless, it will be critical to have additional such studies in order to assess the degree to which the patterns of FC reported here can be generalized across different stroke and participant characteristics – critically lesion location and size, sex, education level, among others. This study involved only left hemisphere stroke and it will be especially important to understand if there are differential consequences depending on the hemisphere of stroke. Siegel et al (2016,2018) included both left and right hemisphere strokes, but the FC analyses did not distinguish between them. A second major limitation is the absence of behavioral data with which to evaluate the consequences of the FC changes observed. As a result, we cannot understand which of the FC patterns observed are associated with positive behavioral outcomes and, if so, in which cognitive domains. This type of information may be particularly useful in making intervention-related decisions, including neurostimulation options.

## 5. Conclusions

This longitudinal investigation of changes in functional connectivity networks specifically examined changes in modularity, hemispheric symmetry and inter-hemispheric communication over the first year following a left hemisphere stroke. It reveals a complex picture of the dynamic changes the brain undergoes as it responds to abrupt onset damage and reorganizes in an attempt to optimize cognitive functioning, resulting in changes that may be adaptive and optimal, but could also be maladaptive and suboptimal.

## Acknowledgements

We gratefully acknowledge and thank our research subjects and their caregivers for their willingness to participate in this research. We thank Jolie Anderson, Miranda Brenneman, Cris Hamilton, Danielle Rossi, and Chia-Ming Lei for data collection. We thank Cris Hamilton for providing lesion mask demarcation. We thank the clinical neurological intensive care unit teams at the University of Texas Health Houston and the Memorial Hermann Hospital System – Texas Medical Center, The Houston Methodist Hospital and Research Institute, and the Baylor St. Luke’s Hospital for their assistance in patient recruitment and neurological assessment.

## Footnotes

1 Formula of weighted *modularity*: 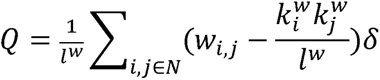 *L^w^* : *sum of all weights in the network* *w_i,j_* : *connection weight between node i and j* 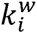 : *weighted degree of node i* *δ_i,j_*: *δ* = 1 *if i and j are in the same network, otherwise δ* = 0

