## Supplementary material for "Longitudinal changes in functional connectivity networks in the first year following stroke": Table S1

**Supplementary Table 1. Hemispheric *modularity* results of the subgroup (n=7).** Longitudinal comparisons were conducted by chronological order, i.e., positive t-value indicates increase over time and vice versa. For comparisons to controls, positive-t-values indicates stroke group higher than controls and negative t-values indicates stroke group lower than controls.

**Left hemisphere**

|  | t-value | p-value |
| --- | --- | --- |
| *Longitudinal changes* | | |
| 1 – 6 month | -0.389 | 0.704 |
| 6 – 12 month | -1.485 | 0.163 |
| 1 – 12 month | -1.878 | 0.085 |
| *Comparisons to healthy controls (HC)* | | |
| 1 month vs. HC | 0.397 | 0.695 |
| 6 month vs. HC | 0.027 | 0.978 |
| 12 month vs. HC | -1.348 | 0.192 |

**Right hemisphere**

|  | t-value | p-value |
| --- | --- | --- |
| *Longitudinal changes* | | |
| 1 – 6 month | -0.165 | 0.871 |
| 6 – 12 month | -2.312 | 0.035 |
| 1 – 12 month | -2.481 | 0.025 |
| *Comparisons to healthy controls (HC)* | | |
| 1 month vs. HC | 0.750 | 0.461 |
| 6 month vs. HC | 0.573 | 0.572 |
| 12 month vs. HC | -1.477 | 0.153 |
